# Hdac3, Setdb1, and Kap1 mark H3K9me3/H3K14ac bivalent regions in young and aged liver

**DOI:** 10.1101/623975

**Authors:** Andrew J. Price, Mohan C. Manjegowda, Irina M. Bochkis

## Abstract

Post-translational modifications of histone tails play a crucial role in gene regulation. Here, we performed chromatin profiling by quantitative targeted mass spectrometry to assess all possible modifications of the core histones. We discovered a novel bivalent combination, a dually-marked H3K9me3/H3K14ac modification in the liver, that is significantly decreased in old hepatocytes. Subsequent genome-wide location analysis (ChIP-Seq) identified 1032 and 668 bivalent regions in young and old livers, respectively, with 280 in common. Histone H3K9 deacetylase Hdac3, as well as H3K9 methyltransferase Setdb1, found in complex Kap1, occupied bivalent regions in both young and old livers, correlating to presence of H3K9me3. Expression of genes associated with bivalent regions in young liver, including those regulating cholesterol secretion and triglyceride synthesis, is upregulated in old liver once the bivalency is lost. Hence, H3K9me3/H3K14ac dually-marked regions define a poised inactive state that is resolved with loss of one or both of the chromatin marks, which subsequently leads to change in gene expression.

## Introduction

Post-translational modifications of histone tails play a crucial role in gene regulation (Jenuwein and Allis, 2001). In particular, tri-methylation of histone H3 lysine 9 (H3K9) is associated with heterochromatin formation (Saksouk et al., 2015) while acetylation of histone H3 lysine 14 (H3K14) is critical for DNA damage and checkpoint activation and circadian regulation (Wang et al., 2012) (Tasselli and Chua, 2015). Typically, acetylation of H3K9 and H3K14 co-occur together, leading to gene activation (Karmodiya et al., 2012). However, levels H3K9me3 and H3K14ac increased together in livers of offspring of mothers fed high fat diet (Suter et al., 2014), leading to metabolic changes.

A specific histone modification pattern of two co-occurring marks, one activating and the other repressing gene expression, named “bivalent domain” was first described in mESCs (Bernstein et al., 2006). These dually marked H3K4me3/H3K27me3 regions were in a poised inactive state, with one mark transmitted to one cell type, leading to gene activation, and the other to another cell type, leading to gene repression, upon differentiation. The role of bivalent domains in differentiated tissues is less clear. Presence of another bivalent mark, consisting of H3K9me3 and H3K14ac modifications has been recently reported in mESCs where the authors demonstrate that the Tudor domain of Setdb1 recognized H3K14ac, whereas the SET domain of Setdb1 methylated H3K9me3 (Jurkowska et al., 2017). This study linked Setdb1 binding at these bivalent domains to silencing of LINE elements.

We have recently connected epigenetic changes to metabolic alterations in aged liver, finding that both altered nucleosome occupancy and redistribution of lamina-associated domains are correlated with development of hepatic steatosis (Bochkis et al., 2014; Whitton et al., 2018) Since histone modification patterns are altered in many organisms with aging (Sidler et al., 2017), we set out to investigate which chromatin marks were altered in old hepatocytes. Chromatin profiling by quantitative target mass spectroscopy identified a novel bivalent H3K9me3/H3K14ac mark in the liver, which is significantly decreased in old hepatocytes. Subsequent ChIP-Seq analysis identified 1032 and 668 bivalent regions in young and old livers, respectively, with 280 in common. Histone H3K9 deacetylase Hdac3, as well as H3K9 methyltransferase Setdb1, found in complex Kap1, occupied bivalent regions in both young and old livers, correlating to presence of H3K9me3. Expression of genes associated with bivalent regions in young liver, including those regulating cholesterol secretion and triglyceride synthesis, is upregulated in old liver once the bivalency is lost. Hence, H3K9me3/H3K14ac dually-marked regions define a poised inactive state that is resolved with loss of one or both of the chromatin marks, which subsequently leads to change in gene expression.

## Results

### New bivalent chromatin state identified by proteomic analysis in mammalian liver

We have previously reported that changes in nucleosome occupancy are associated with metabolic dysfunction in aged livers (Bochkis et al., 2014). In addition, numerous modifications of histone tails have been altered with aging in many cell types (Sidler et al., 2017). Hence, we decided to investigate all post-translational modifications on histone tails in an unbiased manner to determine which chromatin marks change in aged fatty liver. Chromatin profiling by quantitative targeted mass spectrometry (Creech et al., 2015) targeted both individual and combinations of chromatin modifications that resided on the same histone tail, identifying co-occurrence of H3K9me3 and H3K14ac on histone tails from young (3 months) and old (21 months) mouse livers. Presence of this bivalent modification quantitatively decreased in old livers (Figure 1A). We decided to investigate the dual mark for two reasons. First, changes in heterochromatin, marked by H3K9me3, were implicated to drive human aging (Zhang et al., 2015). And second, hepatic levels of both H3K9me3 and H3K14ac increased in offspring of mothers fed high fat diet (Suter et al., 2014), suggesting these modifications are correlated to metabolic changes.

**Figure 1.**
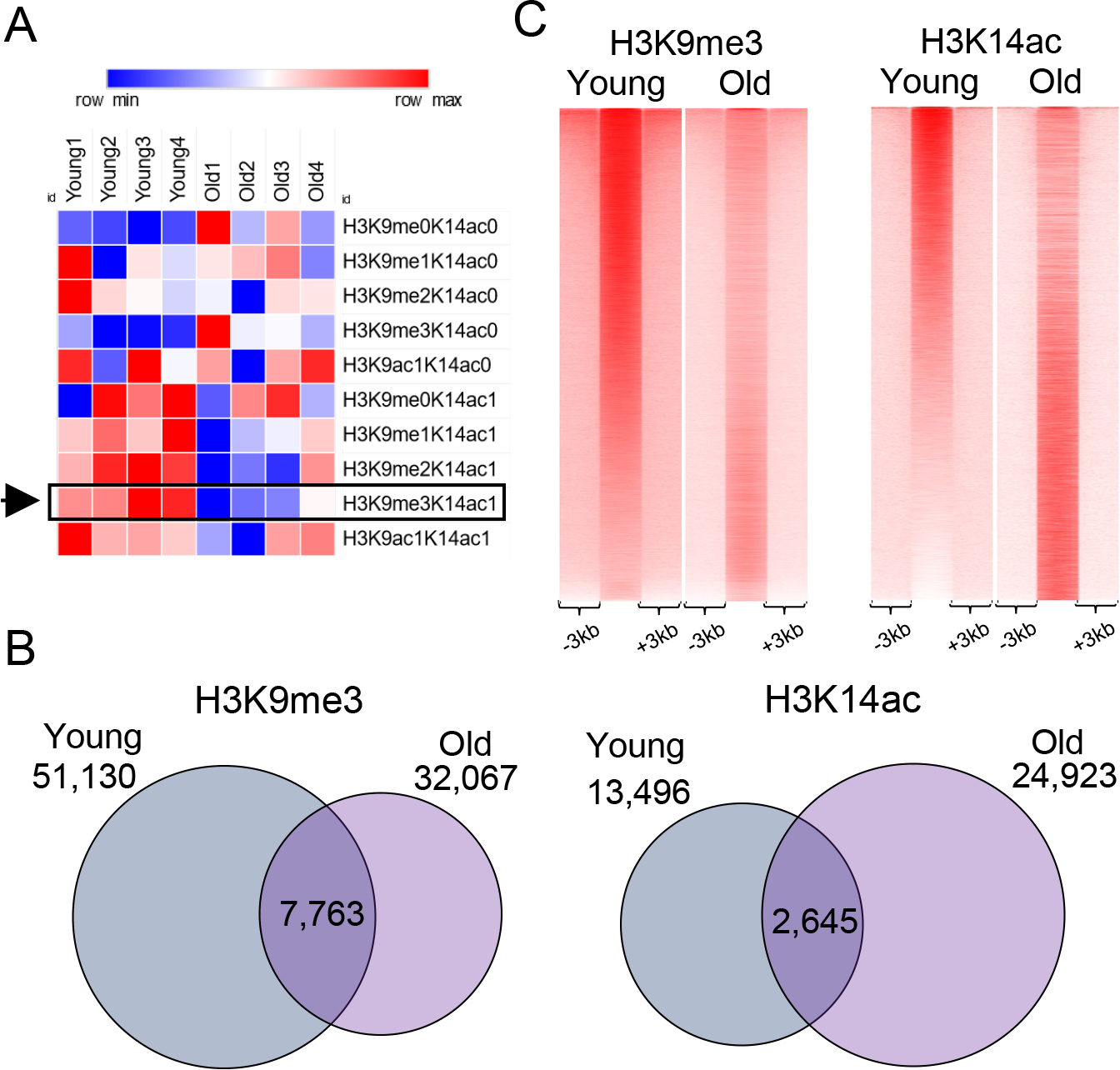
New bivalent chromatin state identified by proteomic analysis in mammalian liver. (**A**) Heatmap showing the relative intensities of histone marks determined by targeted mass-spectrometry of histone tails in young and old livers (values are normalized to row’s median). New bivalent H3K9me3/K14ac mark (marked with a rectangle and an arrow) was identified in both young and old livers and was quantitatively reduced in old livers. (**B**) Venn diagram showing the results of genome-wide location analysis for H3K9me3 (left panel) and H3K14ac (right panel) marks in young and old liver. For H3K9me3, Peak-Seq identified 51,130 sites in young liver and 32,0676 in old liver, with 7,763 in the overlap. For H3K14ac, 13,496 peaks in young liver and 24,923 peaks in old liver were called, with 2,645 in the overlap. (**C**) Heatmaps comparing H3K9me3 (left panel) and H3K14ac (right panel) ChIP-Seq coverage in young and old livers. Overall, H3\K9me3 mark is reduced while H3K14ac modification is increased in old livers. Reads were merged from two replicates for H3K9me3 and H3K14ac ChIP-Seq data in both conditions. Data for H3K9me3 were accessed from our previous study (Whitton et al., 2018).

Next, we performed genome-wide location analysis (ChIP-Seq) of H3K9me3 (previous study (Whitton et al., 2018)) and H3K14ac to determine the genomic regions marked by the dual modification. We identified 51,130 peaks in young and 32,067 peaks in the old livers marked by H3K9me3, with an overlap of 7,763. Similar analysis established 13,496 peaks in young and 22,363 peaks in old livers marked by H3K14ac, with an overlap of 2,645 (Figure 1B). Overall, there was more H3K9me3 but less H3K14ac modification in young livers as compared to the old (Figure 1C). Next, comparing binding regions for both marks, we identified 1032 bivalent sites in young and 668 in old livers, with 280 in common (Figure 2A, examples in 2B). This is consistent with mass spectroscopy results that showed quantitative decrease in bivalent modification on histone tails in old livers (Figure 1A).

**Figure 2.**
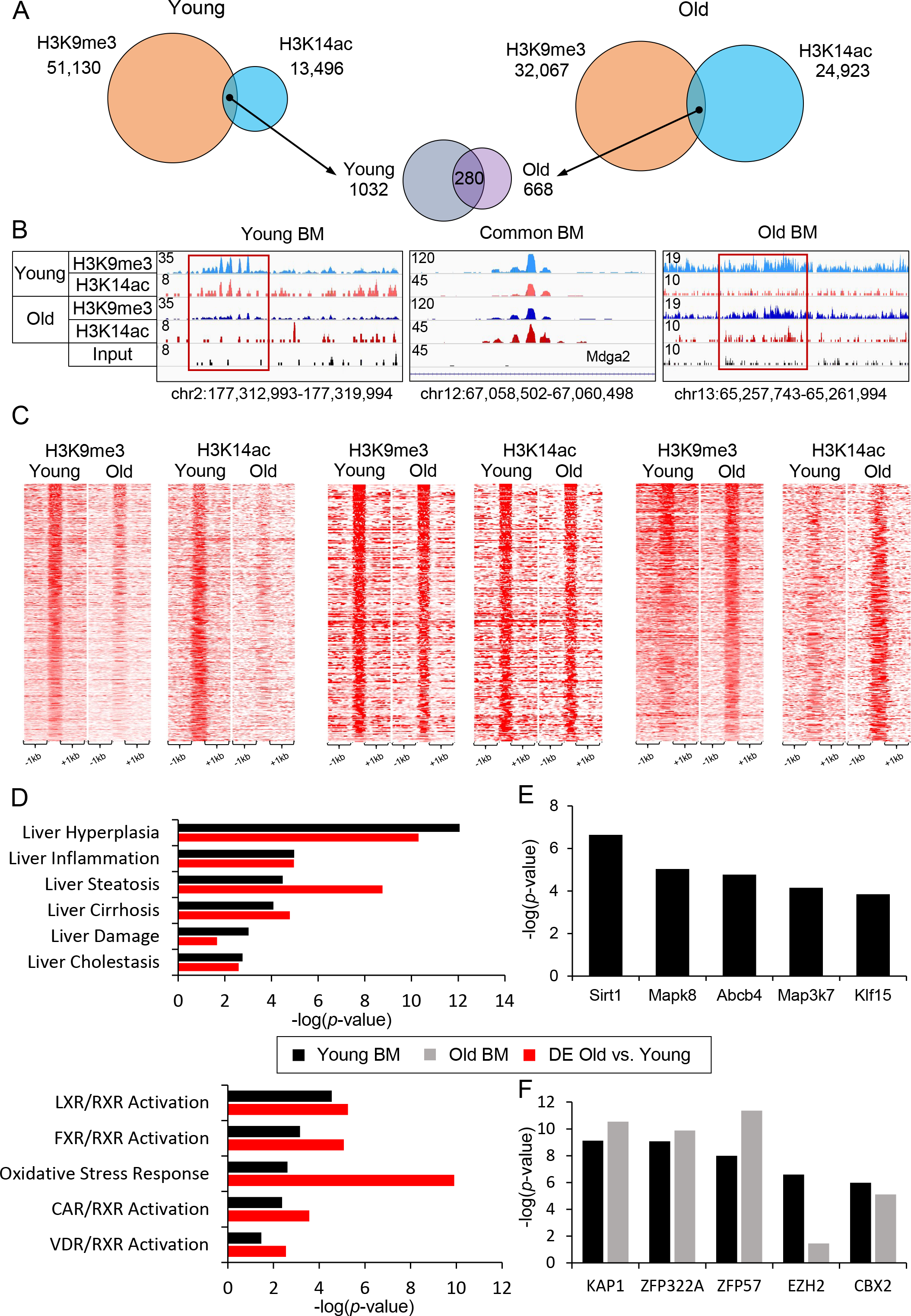
Bivalent regions in young are distinct from those of aged liver. (**A**) Venn diagrams showing the overlap of H3K9me3 and H3K14ac marks in young (left panel) and old (right panel) livers, identifying 1032 and 668 bivalent regions (H3K9me3/K14ac) with 280 domains were present at both conditions (middle panel). (**B**) Examples of genomic regions with H3K9me3/H3K14ac bivalent mark specific to young liver (left panel: chr2:177,312,993-177,319,994, bivalent region in red rectangle), common to both (middle panel: chr12:67,058,502-67,060,498). and old livers (right panel: chr13:65,257,743-65,261,994, bivalent region in red rectangle). Magnitude of the ChIP-Seq signal is shown on y-axis. For comparison, tracks of a given mark in young and old conditions are group scaled and input track is set to the lowest of the magnitudes in the view. (**C**) Heatmaps showing H3K9me3 and H3K14ac ChIP-Seq signal at bivalent regions specific to young (left panel), common to both (middle panel), and specific to old livers (right panel). (**D**) Comparison of over-represented disease functions (top bar graph) and pathways (bottom bar graph) identified by Ingenuity Pathway Analysis of genes associated with bivalent mark in young livers (Young BM black bar) and differentially regulated genes in aged livers (DE Old vs. Young, red bar). Genes associated with bivalent regions are identified by Genomic Regions Enrichment of Annotations Tool (GREAT, see methods for details). (**E**) Single gene perturbation analysis by Enrichr calculates statistical significance of the overlap of a gene set differentially expressed in a knockout of each factor and the input gene set. Analysis of enriched gene sets in genes associated with bivalent mark in young livers (Young BM) identified Sirt1 target genes from livers *Sirt1* KO mice as the most significantly enriched (*p*-value 2.28×10^−07^) among Young BM. (**F**) Chromatin-x Enrichment Analysis (ChEA 2016) of bivalent regions of young (Young BM, black bar) and old (Old BM, gray bar) livers by Enrichr shows Kap1 (*p*-value young: 2.89×10^−11^, old: 7.55×10^−10^) binding sites are most significantly enriched at these bivalent regions. Reads are merged from two replicates in each condition. ChIP-Seq data for H3K9me3 (Whitton et al., 2018) and RNA-Seq data (Bochkis et al., 2014) for differentially regulated genes in aged liver are from our previous studies.

Bivalent regions specific to young livers lose both marks in old hepatocytes. In contrast, some bivalent regions specific to old livers are also marked by H3K9me3 in young hepatocytes. The key difference is absence of H3K14ac mark in young livers and gain of the mark, creating bivalent sites in old hepatocytes (Figure 2C). Next, we mapped regions with bivalent marks to closest genes using ChIP-Seq GREAT (McLean et al., 2010) and compared them to differentially expressed transcripts in old livers (RNA-Seq) (Bochkis et al., 2014) using Ingenuity Pathway Analysis. There are many similarities between genes residing in dually-marked regions in young livers and genes whose expression changes in old livers, with nuclear receptor (LXR, FXR, CAR, and VDR) activation pathways comparably enriched (Figure 2D). This comparison suggests that genes in H3K9me3/H3K14ac bivalent regions are in poised inactive state in adult liver, similar to classical H3K4me3/H3K27me3 bivalent domains characterized in mESCs (Bernstein et al., 2006). Genes regulated by nuclear receptor gene pathways in bivalent regions in young livers get activated in old hepatocytes.

In contrast, only several pathways are similar between genes mapped from dually-marked regions in old livers and differentially expressed genes (“Oxidative Stress”, p-value < 8.51E-3, “Liver Proliferation”, p-value <8.51E-3, “Ahr signaling”, p-value<9.77E-3). Additional functional analysis using EnrichR (Kuleshov et al., 2016) identified Sirt1 as a regulator of expression of genes associated with bivalent regions in young livers as the overlap between those genes and genes differentially expressed in liver-specific *Sirt1* KO mice (GSE14921) was highly significant (p-value < 2.28E-7, Figure 2E). In addition, Kap1 binding sites (ChIP-Seq, GSE31183) were significantly overrepresented among both regions identified as dually-marked in both young and old livers (p-values < 7.55E-10 and < 2.90E-11, respectively, Figure 2F). Sirt1 deacetylates H3K14ac (Imai et al., 2000), while Kap1 interacts with Setdb1 (Schultz et al., 2002), a chromatin regulator that is associated with H3K9me3/H3K14ac dually-marked regions in mESCs (Jurkowska et al., 2017). Hence, we examined the role of both these regulators in bivalent regions.

### Sirt1 regulates genes associated with bivalently-marked regions

We have shown that while old-specific bivalent regions also exhibit H3K9me3 binding at some regions, absence of H3K14ac mark in young livers and key gain of that mark creates bivalent sites in old hepatocytes (Figure 2C). Sirt1, a H3K14 deacetylase, is implicated to govern gene expression in dually-marked regions by EnrichR analysis (Figure 2E). Hence, we set out to examine the role of Srt1 in dually-marked regions. Expression of Sirt1 decreases in old livers (Figure 3A), consistent with a previous report (Jin et al., 2011), and correlating to increased H3K14 acetylation in aged hepatocytes (Figure 1C). In addition, we propose that deacetylase activity of Sirt1 is important to establishing the bivalent regions. While inhibition of Sirt1 activity correlates with presence of dually-marked regions, activation of Sirt1 activity correlates with loss of bivalency (Figure 3B). Next, we compared genes regulated by Sirt1 in the liver (Sirt1 liver KO, GSE14921) with genes associated with young and old bivalent regions. The overlap consists of 191 and 175 Sirt1-regulated genes residing in dually-marked sites in young and old livers, respectively, with overwhelming majority being downregulated (91% in young and 88% in old livers). While H3K14ac is considered an activating chromatin mark, it has also been shown to mark inactive inducible promoter in mESCs (Karmodiya et al., 2012). Hence, genes downregulated in *Sirt1* KO livers correspond to those present in bivalent regions where gene expression is poised but inactive. This is consistent with our model where lack of Sirt1 activity correlates with presence of bivalent poised region (Figure 3B). Comparison of Sirt1-regulated genes associated with bivalent regions in young liver and differentially expressed transcripts in old livers by IPA identified common pathways including “LXR activation”, “Mitochondrial Function”, and “Liver Steatosis” (p-values < 5.49E-6, 8.12<−4, and 2.0<−9, Figure 3C). These genes are poised in young livers and activated in old hepatocytes. In contrast, EnrichR analysis of Sirt1-regulated gens associated with dually-marked sites in old liver detected pathways associated with senescence, including formation of senescence-association heterochromatin foci, which contribute to repression of genes promoting proliferation (Narita et al., 2003), consistent with reduction of proliferative capacity in aged livers (Iakova et al., 2003) (Figure 3D).

**Figure 3.**
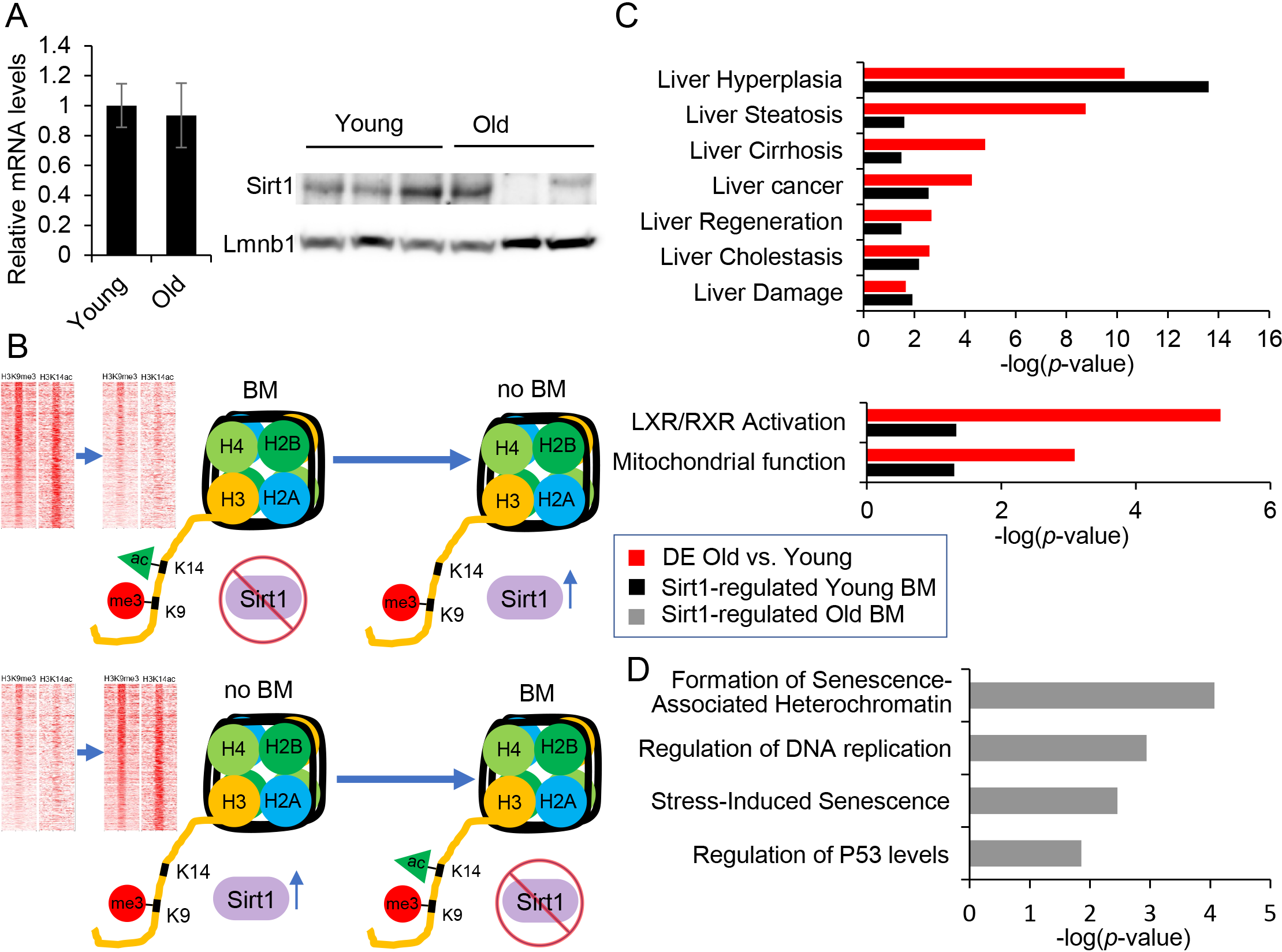
Sirt1 regulates genes associated with bivalently-marked regions. (**A**) Relative mRNA levels (n=4, left panel) of Sirt1 by quantitative RT-PCR. Gapdh was used as a housekeeping gene. Sirt1 mRNA levels do not change in old livers. Western blot analysis (n=3, right panel) of protein nuclear extracts with antibodies to Sirt1 and Lmnb1 (loading control) in young and old livers. Protein levels of Sirt1 are decreased in old livers. (**B**) Model implicating the histone deacetylase, Sirt1, as a key regulator of H3K9me3/K14ac bivalent mark (BM). Sirt1-mediated deacetylation of H3K14ac leads to loss of bivalent mark (no BM, top panel) and loss of Sirt1 activity allows for formation of new domains (BM, bottom panel). In each case, heatmaps show the corresponding changes in the H3K14ac signal. (**C**) Comparison of overrepresented disease functions (top panel) and pathways (bottom panel) identified by Ingenuity Pathway Analysis between differentially expressed genes in aged liver (DE Young vs. Old, red bar) and Sirt1-regulated genes associated with young liver bivalent mark (Sirt1-regulated Young BM, black bar). (**D**) Analysis of over represented pathways in Sirt1-regulated genes associated with old liver-specific bivalent regions by Enrichr (Sirt1-regulated Old BM, gray bar). Formation of senescence associated heterochromatin (*p*-value 8.56×10^−5^) was most significantly enriched pathway. Data set for differentially regulated genes in aged livers is an RNA-Seq data set from our previous study (Wang et al., 2012) and for Sirt1-regulated genes is from a published microarray study (Purushotam et al., 2009).

### Binding of Hdac3, Setdb1, and Kap1 associated with bivalent regions

We have previously implicated Hdac3, a H3K9 deacetylase, with a role in age-associated hepatic steatosis (Bochkis et al., 2014). We hypothesized that Hdac3 activity could be important in establishing the dually-marked sites. Strikingly, Hdac3 binding sites (ChIP-Seq, GSE60393 from (Bochkis et al., 2014)) correlated with presence of H3K9me3 signal in about half of bivalent regions in both young and old livers (Figure 4A). We find that Hdac3 binds bivalent regions that are specific to young livers and common to young and old livers but is not occupying old-specific bivalent regions. Examples of Hdac3-bound bivalent regions include apolipoprotein genes in young and inflammatory genes in old livers (Figure 4B). We propose a model where first a histone acetyltransferase acetylates both H3K9 and H3K14, known to be acetylated together in many regions (Karmodiya et al., 2012), followed by deacetylation of H3K9 residue by Hdac3 allowing for its subsequent tri-methylation (Figure 4C). Next, in order to differentiate between bivalent regions bound versus not occupied by Hdac3, we performed motif scanning analysis using PscanChIP (Zambelli et al., 2013). Interestingly, Onecut/HNF6 motif was highly enriched in Hdac3-bound regions in both young and old livers (p-values < 3.1E-23 and 2.2E-16, respectively, Figure 4D), suggesting that Hdac3 is binding DNA in complex with HNF6 at these bivalent sites. Our analysis is consistent with a previous report implicating HNF6/Hdac3 complex in regulation of hepatic lipid metabolism (Zhang et al., 2016). In contrast, motifs in dually-marked regions without Hdac3 binding included AP-2 sequence (p-value < 0.0098) and consensus for estrogen receptor (ER, p-value< 0.0095) in young livers and E-boxes bound by bHLH factors in old livers (p-values <0.0069 and <0.0118, respectively).

**Figure 4.**
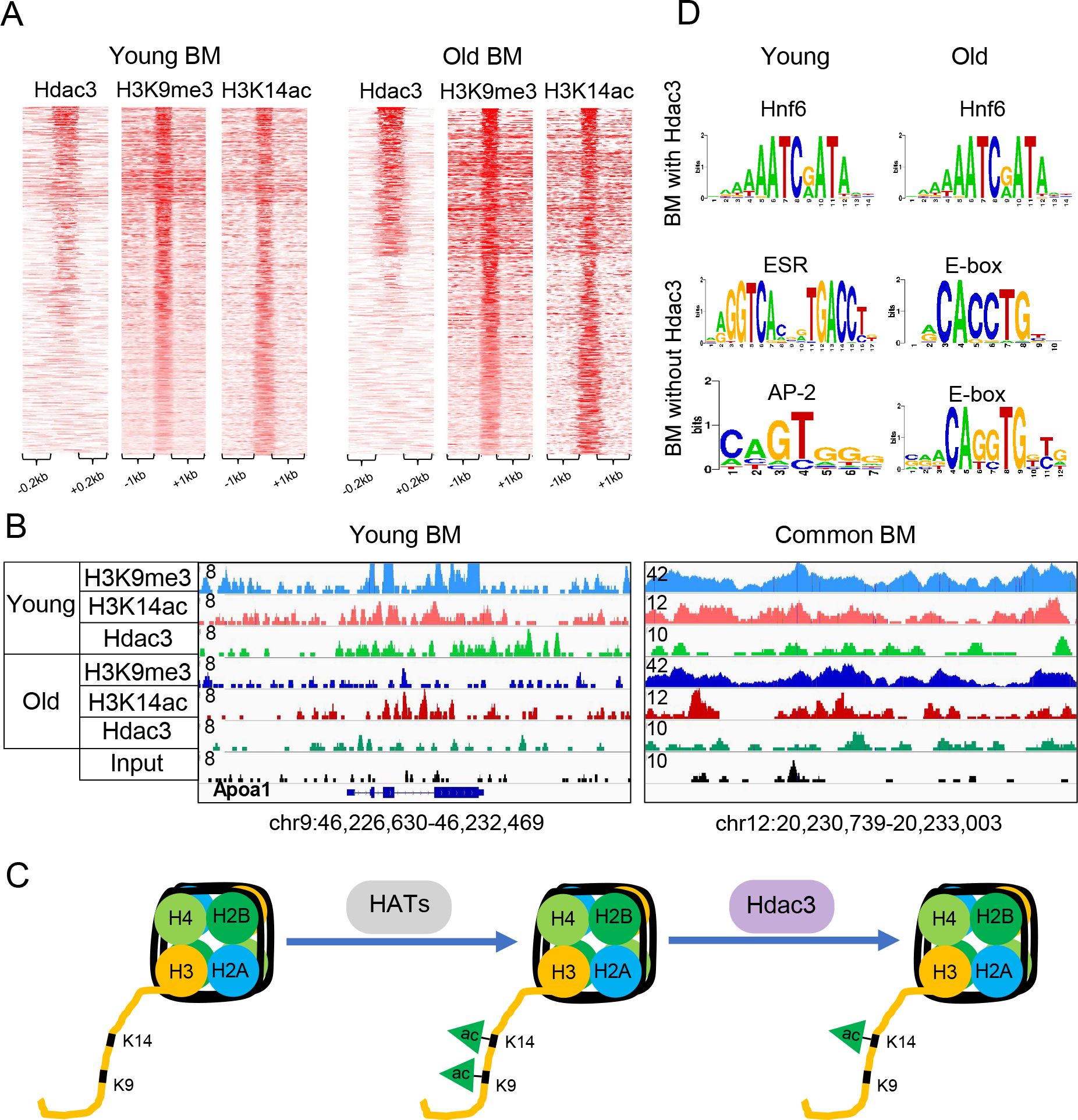
Hdac3 occupies H3K9me3/H3K14ac bivalent regions. (**A**) Heatmaps showing Hdac3, H3K9me3 and H3K14ac ChIP-Seq signal at bivalent regions of young (left panel) and old (right panel) livers. A subset of bivalent regions is bound by Hdac3 in both young and old livers. (**B**) Examples of genomic regions showing the bivalent region bound by Hdac3 in young (chr9: 46,226,630-46,232,469) and old (chr12:20,230,739-20,233,003) livers. Magnitude of the ChIP-Seq signal is shown on y-axis. For comparison, tracks of a given mark in young and old conditions are group scaled and input track is set to the lowest of the magnitudes in the view. (**C**) Model implicating histone deacetylase, Hdac3, in establishment of bivalent mark. First, a histone acetyltransferase acetylates both H3K9 and H3K14, known to be acetylated together in many regions, followed by deacetylation of H3K9 residue by Hdac3, allowing for its subsequent tri-methylation. (**D**) Overrepresented motifs in the bivalent regions with and without Hdac3 in young and old livers, identified by PscanChIP. Onecut/Hnf6 motifs were enriched in both young and old bivalent regions bound with Hdac3 (p-values < 3.1E-23 and 2.2E-16). Whereas, bivalent regions not bound with Hdac3 were enriched with consensus for estrogen receptor (ER, p-value< 0.0095) and its co-activator AP-2 (p-value < 0.0098) in young livers and E-boxes bound by bHLH factors in old livers (p-values <0.0069 and <0.0118, respectively). ChIP-Seq data for Hdac3 (Bochkis et al., 2014) and H3K9me3 (Whitton et al., 2018) are from our previous studies.

To complete the assembly of bivalent sites, we propose that deacetylation of H3K9 by Hdac3 is followed by writing of H3K9 triple methylation by a histone methyltransferase Setdb1 (Figure 5A). We identified the overlap of dually-marked regions with Kap1 binding sites to be highly significant in both young and old livers (p-value < 7.55E-10, Figure 2F). Kap1 is often found in complex with Setdb1 (Schultz et al., 2002), a H3K9 methyltransferase associated with H3K14ac/H3K9me3 dual sites in mESCs (Jurkowska et al., 2017). To test the hypothesis, we performed genome-wide location analysis (ChIP-Seq) of Setdb1 and Kap1 binding in young and livers. Setdb1 and Kap1 bind a similar number of sites (51,896 and 48,915, respectively) with an overlap of 2,741 regions (Figure 5B, top panel). Setdb1 binding is dramatically reduced to just 2,781 sites and Kap1 occupancy is moderately decreased to 22,841regions in old livers (Figure 5B, bottom panel). Interestingly, overlap of Setdb1 and Kap1 binding (24%) increases in old livers. Expression of both factors does not change either on mRNA or protein level (Figure 5C). Hence, observed redistribution of binding of these regulators is independent of expression. Since Setdb1 is associated with H3kK9me3/H3K1ac dually-marked regions in mESCs (Jurkowska et al., 2017), we looked for presence of Setdb1/Kap1 complex in bivalent regions in young and old livers. Heatmaps of ChIP-Seq signal show 30% overlap in young (311/1032) and 27% in old (179/668) hepatocytes (Figure 5D).We find that Setdb1/Kap1 complex binds bivalent regions that are specific to young livers and common to young and old livers but is not occupying old-specific bivalent regions (Figure 5E). Similar to bivalent regions bound by Hdac3, PscanChIP analysis identified Onecut/HNF6 motif to be significantly enriched in Setdb1/Kap1-bound bivalent sites in young and old livers (p-values < 2.0E-30 and < 2.2E-16, respectively, Figure 5F). In contrast, motifs in dually-marked regions without Setdb1/Kap1 binding included AP-2 sequence (p-value < 0.0088) and consensus for estrogen receptor (ER, p-value< 0.0057) in young livers. We did not identify any significant motifs in Setdb1/Kap1-bound bivalent regions in old livers.

**Figure 5.**
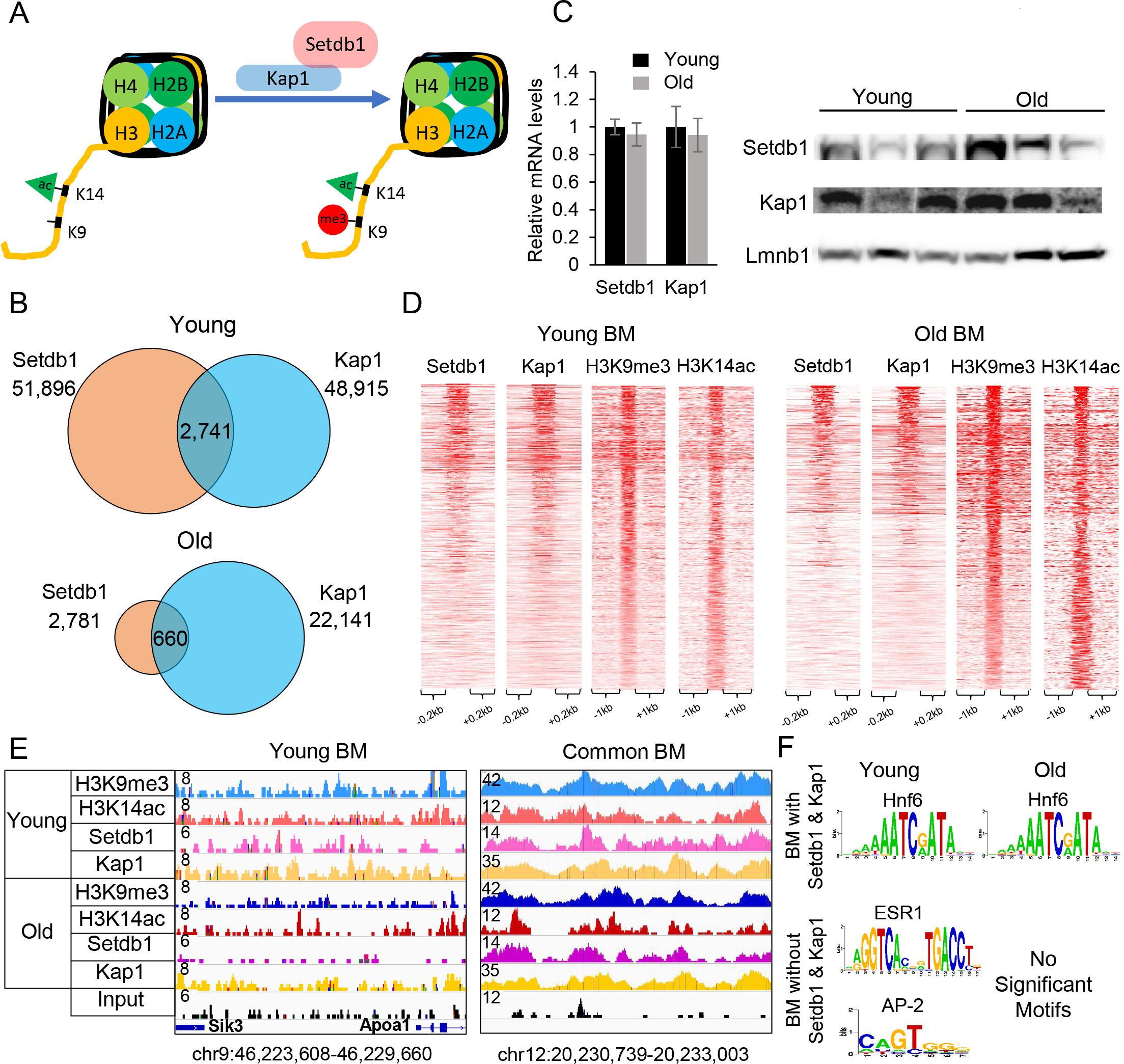
Setdb1/Kap1 complex associates with bivalent regions. (**A**) Model implicating histone methyltransferase, Setdb1, in establishing regions with bivalent mark. Setdb1, in complex with Kap1, methylates H3K9 by recognizing H3K14 acetylation, creating a new dually-marked region. (**B**) Venn diagrams showing the overlap of Setdb1 and Kap1 ChIP-Seq binding sites, identifying 51,896 and 2,781 Setdb1 sites, 48,815 and 22,841 Kap1 sites correspondingly in young (top panel) and old (bottom panel) livers. Of these, 2,741 (young) and 660 (old) sites were bound by both Setdb1 and Kap1 as called by PeakSeq. Reads were merged from two biological replicates in each condition. (**C**) Relative mRNA levels (n=4, left panel) of Setdb1 and Kap1 by quantitative RT-PCR. Western blot analysis (n=3, right panel) of protein nuclear extracts with antibodies to Setdb1, Kap1, and Lmnb1 (loading control) in young and old livers. mRNA and protein expression levels of Setdb1 and Kap1 did not change significantly. Gapdh was used as normalizing control in quantitative RT-PCR. (**D**) Heatmaps showing Setdb1, Kap1, H3K9me3 and H3K14ac ChIP-Seq signal at bivalent regions of young (left panel) and old (right panel) livers. A subset of regions with bivalent mark is bound by Setdb1 and Kap1 in both young and old livers. (**E**) Examples of genomic regions showing bivalent regions bound by Setdb1/Kap1 complex in young (chr9:46,223,608-46,229,660) and common to young and old livers (chr12:20,230,739-20,233,003) livers. Magnitude of the ChIP-Seq signal is shown on y-axis. For comparison, tracks of a given mark in young and old conditions are group scaled and input track is set to the lowest of the magnitudes in the view. (**F**) Overrepresented motifs in the bivalent regions with and without Setdb1/Kap1 binding in young and old livers, identified by PscanChIP. Hnf6 motifs were enriched in bivalent regions occupied by Setdb1/Kap1 in both young and old livers (p-values < 2.0E-30 and < 2.2E-16, respectively). In contrast, motifs in dually-marked regions without Setdb1/Kap1 binding included AP-2 sequence (p-value < 0.0088) and consensus for estrogen receptor (ER, p-value< 0.0057) in young livers. We did not identify any significant motifs in Setdb1/Kap1-bound bivalent regions in old livers. ChIP-Seq data for H3K9me3 are from our previous study (Whitton et al., 2018).

### Hdac3, Setdb1, and Kap1 co-localize to same bivalent regions

Considering that both Hdac3-bound and Setdb1/Kap1 bound bivalent regions were enriched for the sample Onecut/HNF6 motif, we next investigated the possibility if Hdac3 and Setdb1/Kap1 complex were occupying the same sites. Remarkably, the three factors were bound in same regions in both young and old livers (Figure 6A).Our observations are consistent with a previous report showing that N-Cor complex containing Hdac3 interacts with Kap1 (Underhill et al., 2000). Next, we wanted to ascertain whether HNF6 was indeed bound in bivalent regions as motif analysis suggested. We compared HNF6 binding in young liver from a previous study (ArrayExpress E-MTAB-2060)(Wang et al., 2014) finding that occupies a subset of Hdac3/Setdb1/Kap1 bound bivalent sites (30% overlap, 308/1032 of all bivalent sites in young liver, Figure 6A). Examples of HNF6 bound elements at apolipoprotein genes Apoa1 and Apob are shown in Figure 6B. Subsequent comparison of genes in HNF6-bound bivalent regions in young livers and differentially expressed genes in old livers showed an overlap of pathways, including activation of FXR and LXR-dependent gene expression (p-values < 2.0E-4, 1,6,E-4, IPA, Figure 6C). Further, we compared genes differentially expressed in HNF6, Hdac3, and Kap1 knockout livers with genes whose expression changed in old hepatocytes. Remarkably, the overlap included similar pathways enriched in all three knockout models. These include activation of LXR an PPAR-dependent gene expression, consistent with development of steatosis in aged liver (Figure 6D).

**Figure 6.**
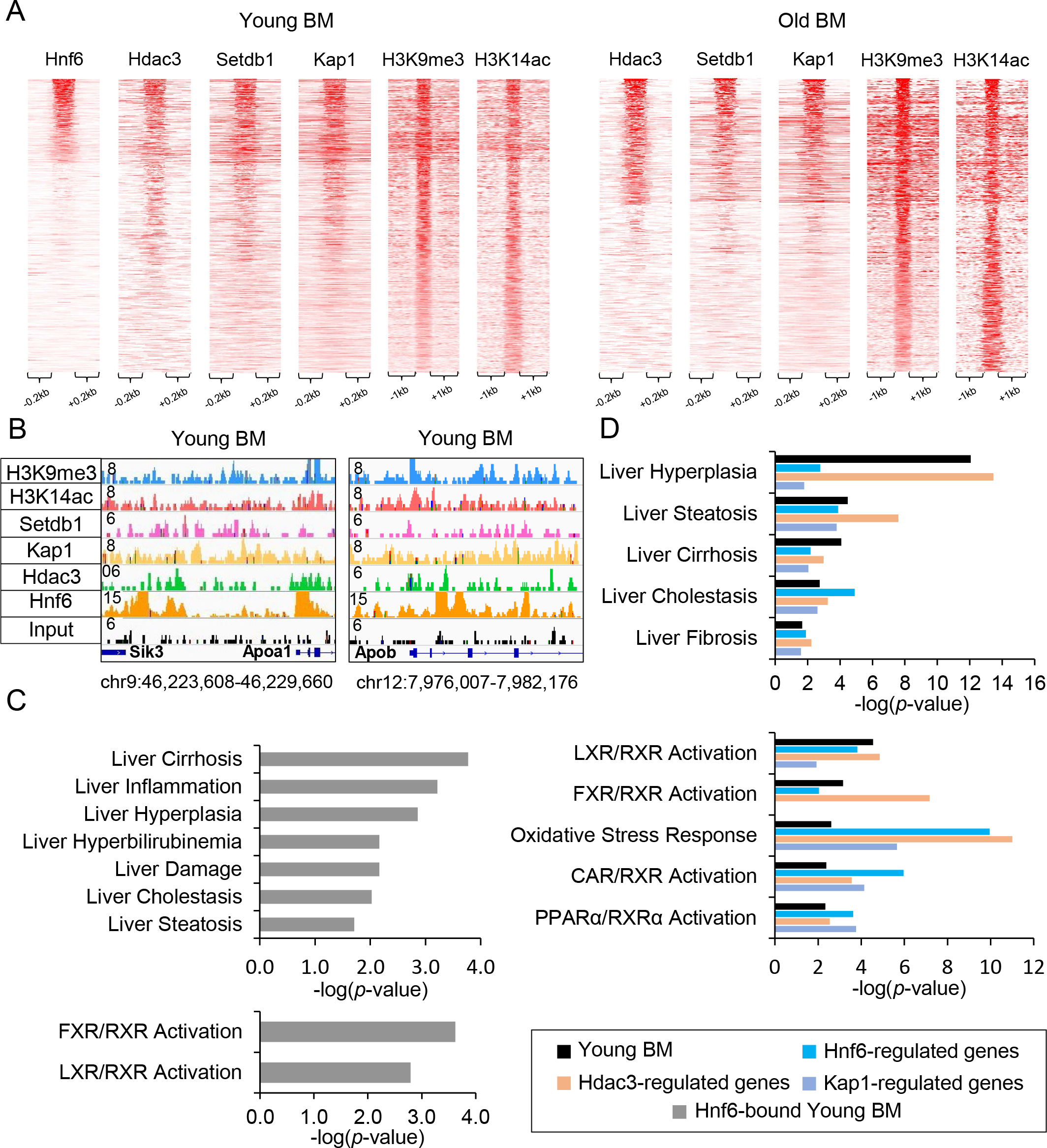
Hdac3, Setdb1, and Kap1 co-localize to same bivalent regions. (**A**) Heatmaps showing Hnf6, Hdac3, Setdb1, Kap1, H3K9me3 and H3K14ac ChIP-Seq signal at regions with bivalent mark in young (left panel) and old livers (right panel). A subset of bivalent regions occupied by both Hdac3 and Setdb1/Kap1 complex is also bound by Hnf6 in young livers. (**B**) Examples of genomic regions showing the bivalent region bound by Hnf6, Hdac3 and Setdb1/Kap1 complex in young livers include regions near two apolipoprotein genes, Apoa1 (chr9:46,223,608-46,229,660) and Apob (chr12:7,976,007-7,982,176). Magnitude of the ChIP-Seq signal is shown on y-axis. For comparison, tracks of a given mark in young and old conditions are group scaled and input track is set to the lowest of the magnitudes in the view. (**C**) Ingenuity Pathway Analysis (IPA) of genes associated with Hnf6-bound bivalent regions in young livers (Hnf6-bound Young BM, gray bar) showing significantly over represented disease functions (top panel) and pathways (bottom panel). Cirrhosis (*p*-value 0.0002), inflammation (*p*-value 0.0006) and hyperplasia (*p*-value 0.0014) are most significantly over represented hepatotoxicity functions, and FXR (*p*-value 0.0002) and LXR (*p*-value 0.0016) activation pathways are most significantly overrepresented. (**D**) Comparison of over represented disease functions (top panel) and pathways (bottom panel) identified by Ingenuity Pathway Analysis (IPA) between young liver bivalent mark-associated genes (Young BM, black bar) and genes regulated by Hnf6 (blue bar), Hdac3 (orange bar) and Kap1 (purple bar). Genes associated with bivalent regions are mapped by Genomic Regions Enrichment of Annotations Tool (GREAT). Expression data for Hnf6 (Zhang et al., 2016), Hdac3 (Sun et al., 2013) and Kap1-regulated (Bojkowska et al., 2012) gene sets are published microarray studies. ChIP-Seq data for Hnf6 are from a published study (Wang et al., 2014) and for H3K9me3 are from our previous study (Bochkis et al., 2014).

In summary, we propose a model where first close residues H3K9 and H3K14 are acetylated by the same histone acetyltransferase (HAT), then H3K9 is deacetylated by Hdac3, allowing for subsequent triple methylation of H3K9 by Setdb1/Kap1 and establishment of the bivalent region (Figure 7). In addition, we propose that Sirt1 activity is inhibited during bivalent region assembly to preserve acetylation of H3K14 that serves as the first step in that process. Our model is consistent with a prior study reporting that, first, established H3K14ac mark is recognized by the Tudor domain of Setdb1, followed by triple methylation of H3K9 by the SET domain of Setdb1 (Jurkowska et al., 2017).

**Figure 7.**
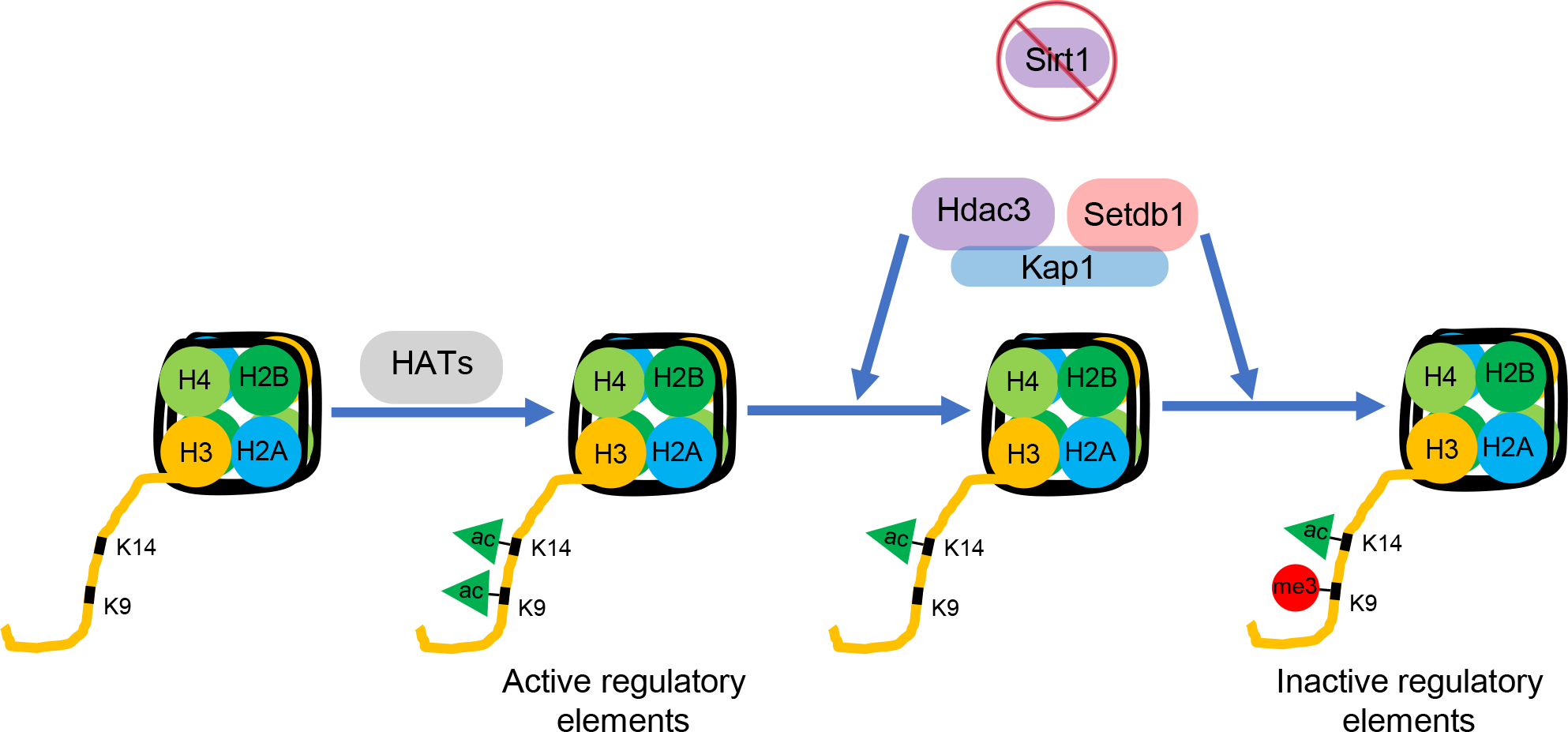
Model describing establishment of H3K9me3/H3K14ac bivalent mark. In the first step, close residues H3K9 and H3K14 are acetylated by the same histone acetyltransferase (HAT). Then, H3K9 is deacetylated by Hdac3, allowing for subsequent triple methylation of H3K9 by Setdb1, which is found in complex with Kap1, and establishment of the bivalent region. Sirt1 activity is inhibited during bivalent region assembly to preserve acetylation of H3K14 that serves as the first step in that process. Activation of Sirt1 activity leads to loss of the bivalent mark.

## Discussion

Here, we performed chromatin profiling by quantitative targeted mass spectrometry to assess all possible modifications of the core histones in young and old liver. We discovered a novel hepatic bivalent combination, a dually-marked H3K9me3/H3K14ac mark, that is significantly decreased in old hepatocytes. Subsequent genome-wide location analysis (ChIP-Seq) identified 1032 and 668 bivalent regions in young and old livers, respectively, with 280 in common. Our ChIP-Seq results were consistent with proteomic analysis showing quantitative decrease of this combination in old livers. Presence of this bivalent mark has been reported previously in mESCs where the authors linked H3K14ac, recognized by the Tudor domain of Setdb1, to H3K9me3, methylated by SET domain of Setdb1 (Jurkowska et al., 2017). The authors reported a larger H3K14ac/H3K9me3 overlap in ES cells (12,400 bivalent regions, 2231 bound by Setdb1). We observe a smaller number of bivalent regions in adult differentiated tissue.

In addition to Setdb1, we also localized Kap1, an interaction partner of Setdb1, and Hdac3, an enzyme that deacetylates H3K9, to bivalent regions. While binding of Hdac3 is typically associated with absence of H3K9ac mark (Feng et al., 2011), we find that Hdac3 correlates with presence of H3K9me3 modification. We propose that Hdac3/Setdb1/Kap1 act in concert, with Hdac3 first deacetylating H3K9, followed by subsequent triple methylation by Setdb1. About half of sites occupied by Hdac3/Setdb1/Kap1 are also bound by HNF6. Hdac3 has been shown to bind DNA in complex with HNF6 (Zhang et al., 2016) at regulatory elements of lipid metabolic genes but a relationship between HNF6 and Setdb1 has not been reported previously. There are several reasons why HNf6 co-localizes with a subset of Hdac3/Setdb1/Kap1 sites. First, Hdac3 binding is circadian (Feng et al., 2011) and a different subset of HNF6-bound regions could overlap with Hdac3 at a different time point. Also, HNF6-bound regions control genes important for hepatic sexually dimorphic gene expression (Conforto et al., 2015) and HNF6 will not occupy other sites bound by Hdac3/Setdb1/Kap1. This possibility is reinforced by presence of motifs for estrogen receptor and its co-activator AP-2 (Tan et al., 2011) in sites not occupied by Hdac3/Setdb1/Kap1, leading to repression of female-specific gene expression in male liver. This observation suggest that H3K9me3/H3K14ac could play a role in regulation of sexually dimorphic gene expression in the liver.

Further, we observe similarities between genes connected to bivalent marks in young liver and those differentially expressed in old liver. Apolipoprotein genes that govern cholesterol export and genes important for triglyceride synthesis are of particular interest. They correspond to gene expression and metabolic changes, such increased cholesterol secretion (Einarsson et al., 1985) and development of hepatic steatosis (Whitton et al., 2018) in old livers. Interestingly, these genes are connected to regions bound by HNF6, Hdac3, and Setdb1/Kap1 and comparison analysis of genes regulated by these factors identified pathways similar to those upregulated in old livers. Together, these observations suggest that H3K9me3/H3K14ac dually-marked regions constitute a poised inactive state in young livers, which is resolved in old livers, leading to changes in gene expression that contribute to metabolic dysfunction.

## Experimental Procedures

### Mice

Young (3 months) and old (21 months) male mice (C57BL/6) were purchased from the National Institute of Aging (NIA) aged rodent colony (Charles River Laboratories). Upon arrival, animals were housed for a week to acclimate to the light-dark cycle at the UVa facility before tissue harvest. Four biological replicates of young and old mice were used for proteomics study. Two biological replicates of young and old mice were used for chromatin immunoprecipitation and sequencing. All animal work was approved by Animal Care and Use Committee at UVa (protocol number 4162-03-17).

### Analysis of histone modifications by targeted Mass Spectrometry

Quantitative targeted Mass-Spectrometry-based profiling of histone modifications was performed at LINCS Proteomic Characterization Center for Signaling and Epigenetics at The Broad Institute of MIT and Harvard University, Cambridge, MA. Detailed experimental procedure for the complete proteomics study is described previously (Creech et al., 2015). Heatmap of relative abundance of histone modifications was drawn by Morpheus (Morpheus, https://software.broadinstitute.org/morpheus).

### RNA and protein analysis

Analysis of mRNA and protein expression levels were performed as described previously (Whitton et al., 2018). Gapdh was used as normalizing gene for quantitative RT-PCR analysis. Primer sequences will be provided upon request. Lmnb1 (Abcam, ab16048) was used as loading control. Rabbit monoclonal antibody for Sirt1 (CST, 9475) and rabbit polyclonal antibodies specific to KAP1 (Abcam, ab10483) and SETDB1 (Proteintech, 11231-1-AP) were used in western blotting at 1:1000 dilution. Student’s two sample *t*-test was used to analyze the Q-PCR data.

### Chromatin immunoprecipitation and sequencing

Snap‐frozen mouse liver (100 mg) from young and old wildtype mice was used to prepare chromatin. ChIP and H3K14ac ChIP-Seq were performed as described previously (Whitton et al., 2018). For Setdb1 and Kap1 a few modifications were incorporated. Particularly, sonication, using Diagenode Bioruptor Pico, was reduced to 11 cycles (30 s pulse on, 30 s pulse off) and libraries were sequenced using Illumina NextSeq 500 following the manufacturer's protocols. Rabbit polyclonal antibodies specific to KAP1 (Abcam, ab10483), SETDB1 (Proteintech, 11231-1-AP), and anti-acetyl-histone-H3 specific to Lys-14 (Millipore, 07-353) were used for immunoprecipitation. Detailed description of *in silico* analysis of sequenced reads is previously described (Whitton et al., 2018). Data from two biological replicates were merged in all ChIP-Seq experiments.

### Functional analysis

Functional analysis of ChIP-Seq peaks were performed as described previously (Whitton et al., 2018) with the following modifications. ChIP-Seq peaks were associated with closest genes by Genomic Regions Enrichment of Annotations Tool (GREAT) with basal plus extension method and distal regions extended up to 100,000 kb (McLean et al., 2010). Overlap of ChIP-Seq peak regions were analyzed by Intervene (Khan and Mathelier, 2017). Heatmaps of ChIP-Seq coverage were generated by deeptools(Ramirez et al., 2014). Expression data from published studies for Sirt1 (GSE14921), Hdac3 (GSE49386), Hnf6 (GSE83789) and Kap1-regulated genes(Bojkowska et al., 2012) were accessed from GEO. Differentially expressed genes from published microarray studies were identified by GEO2R tool(Barrett et al., 2013). Chromatin-x enrichment analysis and analyses for over represented pathways and disease conditions were performed by Enrichr (Kuleshov et al., 2016)

### Accession Numbers

ChIP-Seq data from this study can be accessed at GEO under accession numbers GSEXXX for H3K14ac, Setdb1, and Kap1, GSE60393 for Hdac3, GSE78177 for H3K9me3, respectively, and ArrayExpress experiment E-MTAB2060 for Hnf6.

## Acknowledgements

We thank S. Anakk and T. Harris for critical reading of the manuscript and S. Srabani for taking care of the mice. I.M.B. was supported by National Diabetes and Digesting and Kidney Diseases Institute K01 award DK-101633.

## Author Contributions

A.J.P. analyzed data and developed a new direction for the project, M.C.M. performed experiments and data analysis, I.M.B. developed the project, performed experiments and data analysis, and wrote a draft of the manuscript.

## References

Barrett, T., Wilhite, S.E., Ledoux, P., Evangelista, C., Kim, I.F., Tomashevsky, M., Marshall, K.A., Phillippy, K.H., Sherman, P.M., Holko, M., et al. (2013). NCBI GEO: archive for functional genomics data sets-- update. Nucleic Acids Res 41, D991–995.

Bernstein, B.E., Mikkelsen, T.S., Xie, X., Kamal, M., Huebert, D.J., Cuff, J., Fry, B., Meissner, A., Wernig, M., Plath, K., et al. (2006). A bivalent chromatin structure marks key developmental genes in embryonic stem cells. Cell 125, 315–326.

Bochkis, I.M., Przybylski, D., Chen, J., and Regev, A. (2014). Changes in nucleosome occupancy associated with metabolic alterations in aged mammalian liver. Cell Rep 9, 996–1006.

Bojkowska, K., Aloisio, F., Cassano, M., Kapopoulou, A., Santoni de Sio, F., Zangger, N., Offner, S., Cartoni, C., Thomas, C., Quenneville, S., et al. (2012). Liver-specific ablation of Kruppel-associated box-associated protein 1 in mice leads to male-predominant hepatosteatosis and development of liver adenoma. Hepatology 56, 1279–1290.

Conforto, T.L., Steinhardt, G.F.t., and Waxman, D.J. (2015). Cross Talk Between GH-Regulated Transcription Factors HNF6 and CUX2 in Adult Mouse Liver. Mol Endocrinol 29, 1286–1302.

Creech, A.L., Taylor, J.E., Maier, V.K., Wu, X., Feeney, C.M., Udeshi, N.D., Peach, S.E., Boehm, J.S., Lee, J.T., Carr, S.A., et al. (2015). Building the Connectivity Map of epigenetics: chromatin profiling by quantitative targeted mass spectrometry. Methods (San Diego, Calif.) 72, 57–64.

Einarsson, K., Nilsell, K., Leijd, B., and Angelin, B. (1985). Influence of age on secretion of cholesterol and synthesis of bile acids by the liver. The New England journal of medicine 313, 277–282.

Feng, D., Liu T Fau - Sun, Z., Sun Z Fau - Bugge, A., Bugge A Fau - Mullican, S.E., Mullican Se Fau - Alenghat, T.,Alenghat T Fau - Liu, X.S., Liu Xs Fau - Lazar, M.A., and Lazar, M.A. (2011). A circadian rhythm orchestrated by histone deacetylase 3 controls hepatic lipid metabolism. Science.

Iakova, P., Awad, S.S., and Timchenko, N.A. (2003). Aging reduces proliferative capacities of liver by switching pathways of C/EBPalpha growth arrest. Cell 113, 495–506.

Imai, S., Armstrong, C.M., Kaeberlein, M., and Guarente, L. (2000). Transcriptional silencing and longevity protein Sir2 is an NAD-dependent histone deacetylase. Nature 403, 795–800.

Jenuwein, T., and Allis, C.D. (2001). Translating the histone code. Science 293, 1074–1080.

Jin, J., Iakova, P., Jiang, Y., Medrano, E.E., and Timchenko, N.A. (2011). The reduction of SIRT1 in livers of old mice leads to impaired body homeostasis and to inhibition of liver proliferation. Hepatology 54, 989–998.

Jurkowska, R.Z., Qin, S., Kungulovski, G., Tempel, W., Liu, Y., Bashtrykov, P., Stiefelmaier, J., Jurkowski, T.P., Kudithipudi, S., Weirich, S., et al. (2017). H3K14ac is linked to methylation of H3K9 by the triple Tudor domain of SETDB1. Nature communications 8, 2057.

Karmodiya, K., Krebs, A.R., Oulad-Abdelghani, M., Kimura, H., and Tora, L. (2012). H3K9 and H3K14 acetylation co-occur at many gene regulatory elements, while H3K14ac marks a subset of inactive inducible promoters in mouse embryonic stem cells. BMC genomics 13, 424.

Khan, A., and Mathelier, A. (2017). Intervene: a tool for intersection and visualization of multiple gene or genomic region sets. BMC Bioinformatics 18, 287.

Kuleshov, M.V., Jones, M.R., Rouillard, A.D., Fernandez, N.F., Duan, Q., Wang, Z., Koplev, S., Jenkins, S.L., Jagodnik, K.M., Lachmann, A., et al. (2016). Enrichr: a comprehensive gene set enrichment analysis web server 2016 update. Nucleic Acids Res 44, W90–97.

McLean, C.Y., Bristor, D., Hiller, M., Clarke, S.L., Schaar, B.T., Lowe, C.B., Wenger, A.M., and Bejerano, G. (2010). GREAT improves functional interpretation of cis-regulatory regions. Nature biotechnology 28, 495–501.

Narita, M., Nunez, S., Heard, E., Narita, M., Lin, A.W., Hearn, S.A., Spector, D.L., Hannon, G.J., and Lowe, S.W. (2003). Rb-mediated heterochromatin formation and silencing of E2F target genes during cellular senescence. Cell 113, 703–716.

Ramirez, F., Dundar, F., Diehl, S., Gruning, B.A., and Manke, T. (2014). deepTools: a flexible platform for exploring deep-sequencing data. Nucleic Acids Res 42, W187–191.

Saksouk, N., Simboeck, E., and Dejardin, J. (2015). Constitutive heterochromatin formation and transcription in mammals. Epigenetics & chromatin 8, 3.

Schultz, D.C., Ayyanathan, K., Negorev, D., Maul, G.G., and Rauscher, F.J., 3rd (2002). SETDB1: a novel KAP-1-associated histone H3, lysine 9-specific methyltransferase that contributes to HP1-mediated silencing of euchromatic genes by KRAB zinc-finger proteins. Genes & development 16, 919–932.

Sidler, C., Kovalchuk, O., and Kovalchuk, I.. (2017). Epigenetic Regulation of Cellular Senescence and Aging. Frontiers in genetics 8, 138.

Suter, M.A., Ma, J., Vuguin, P.M., Hartil, K., Fiallo, A., Harris, R.A., Charron, M.J., and Aagaard, K.M. (2014). In utero exposure to a maternal high-fat diet alters the epigenetic histone code in a murine model. American journal of obstetrics and gynecology 210, 463.e461–463.e411.

Tan, S.K., Lin, Z.H., Chang, C.W., Varang, V., Chng, K.R., Pan, Y.F., Yong, E.L., Sung, W.K., and Cheung, E. (2011). AP-2gamma regulates oestrogen receptor-mediated long-range chromatin interaction and gene transcription. The EMBO journal 30, 2569–2581.

Tasselli, L., and Chua, K.F. (2015). Methylation gets into rhythm with NAD(+)-SIRT1. Nat Struct Mol Biol 22, 275–277.

Underhill, C., Qutob, M.S., Yee, S.P., and Torchia, J. (2000). A novel nuclear receptor corepressor complex, N-CoR, contains components of the mammalian SWI/SNF complex and the corepressor KAP-1. J Biol Chem 275, 40463–40470.

Wang, L., Chen, J., Wang, C., Uuskula-Reimand, L., Chen, K., Medina-Rivera, A., Young, E.J., Zimmermann, M.T., Yan, H., Sun, Z., et al. (2014). MACE: model based analysis of ChIP-exo. Nucleic Acids Res 42, e156.

Wang, Y., Kallgren, S.P., Reddy, B.D., Kuntz, K., Lopez-Maury, L., Thompson, J., Watt, S., Ma, C., Hou, H., Shi, Y., et al. (2012). Histone H3 lysine 14 acetylation is required for activation of a DNA damage checkpoint in fission yeast. J Biol Chem 287, 4386–4393.

Whitton, H., Singh, L.N., Patrick, M.A., Price, A.J., Osorio, F.G., Lopez-Otin, C., and Bochkis, I.M. (2018). Changes at the nuclear lamina alter binding of pioneer factor Foxa2 in aged liver. Aging Cell 17, e12742.

Zambelli, F., Pesole, G., and Pavesi, G. (2013). PscanChIP: Finding over-represented transcription factor-binding site motifs and their correlations in sequences from ChIP-Seq experiments. Nucleic Acids Res 41, W535–543.

Zhang, W., Li, J., Suzuki, K., Qu, J., Wang, P., Zhou, J., Liu, X., Ren, R., Xu, X., Ocampo, A., et al. (2015). Aging stem cells. A Werner syndrome stem cell model unveils heterochromatin alterations as a driver of human aging. Science 348, 1160–1163.

Zhang, Y., Fang, B., Damle, M., Guan, D., Li, Z., Kim, Y.H., Gannon, M., and Lazar, M.A. (2016). HNF6 and Rev-erbalpha integrate hepatic lipid metabolism by overlapping and distinct transcriptional mechanisms. Genes and Development.

